# Perceived direction of deformation flow

**DOI:** 10.1101/693762

**Authors:** Takahiro Kawabe

**Affiliations:** NTT Communication Science Laboratories

## Abstract

In everyday circumstances, human observers can easily discriminate the direction of transparent liquid flow. However, the mechanism of direction discrimination is not so straightforward. The present study focused on the flow of image deformation, which is closely related to the flow of transparent liquid in the natural world. To determine what image information is important in discriminating the direction of deformation flow, a natural image in a stimulus clip was deformed by using a deformation vector map that translated leftward or rightward. The task of the observers was to judge whether the transparent liquid in the clip flowed leftward or rightward. Manipulating the amplitude of deformation, we found that the discrimination performance improved with the amplitude. Interestingly, the observers’ performance was high overall only when shearing deformation was applied to the stimuli, while the observers reported an opposite-motion direction when only compressive deformation was applied. We computationally analyzed motion statistics of stimuli and found that the combination of mean and skewness of horizontal motion vectors reliably predicted the performance. The results indicate that human observers use global motion directions in order to determine the direction of deformation flow.

## Introduction

Human observers are sensitive to spatiotemporal flow. For example, it is well known that the visual system has detectors dedicated to spatiotemporal luminance flow [1,2]. In addition, the visual system is sensitive to image flow that is defined by visual features other than luminance [3]. For example, human observers can discriminate the direction of contrast-defined flow, though attention is necessary for precise direction discrimination [4,5].

Recently, stimuli related to a new class of image flow, that is, deformation flow, have been investigated. A previous study [6] showed that deformation flow was a source of transparent liquid perception. One study has shown that the deformation flow was perceived to come from transparent liquid when the amplitude of image deformation was large, while it was perceived to come from transparent hot air when the amplitude was small [7]. The local image motion in the deformation flow is also an important determinant of transparent liquid perception: less linear motion coverage in the deformation flow leads to a stronger impression of a transparent liquid flow [8]. In this way, these previous studies have successfully shown that dynamic image deformation was a cue to transparent material perception.

On the other hand, no studies have addressed the following question: how do observers discriminate the direction of flow that is defined by image deformation? At a glance, the deformation flow would seem to be defined by the sequential image deformation of a background image. The visual system may thus be able to monitor the spatiotemporal flow of image deformation and discriminate its direction.

To anticipate key features in the deformation flow, it is useful to discuss elementary deformation related to deformation flow. A previous study [6] found that the flow could be decomposed into two elemental motion vectors, that is, horizontal and vertical deformation (Figure 1). Given a transparent liquid flowing horizontally, the elemental horizontal and vertical deformations can be rephrased as compressive and shearing deformations, respectively. Some previous studies have shown that the visual system has detectors for shear and compressive motion patterns [9], which are likely involved with shearing [10] and compressive deformations, respectively. Thus, it was expected that the visual system discriminated the direction of the deformation flow by checking the spatiotemporal variation of the compressive and/or shearing deformations.

**Figure 1.**
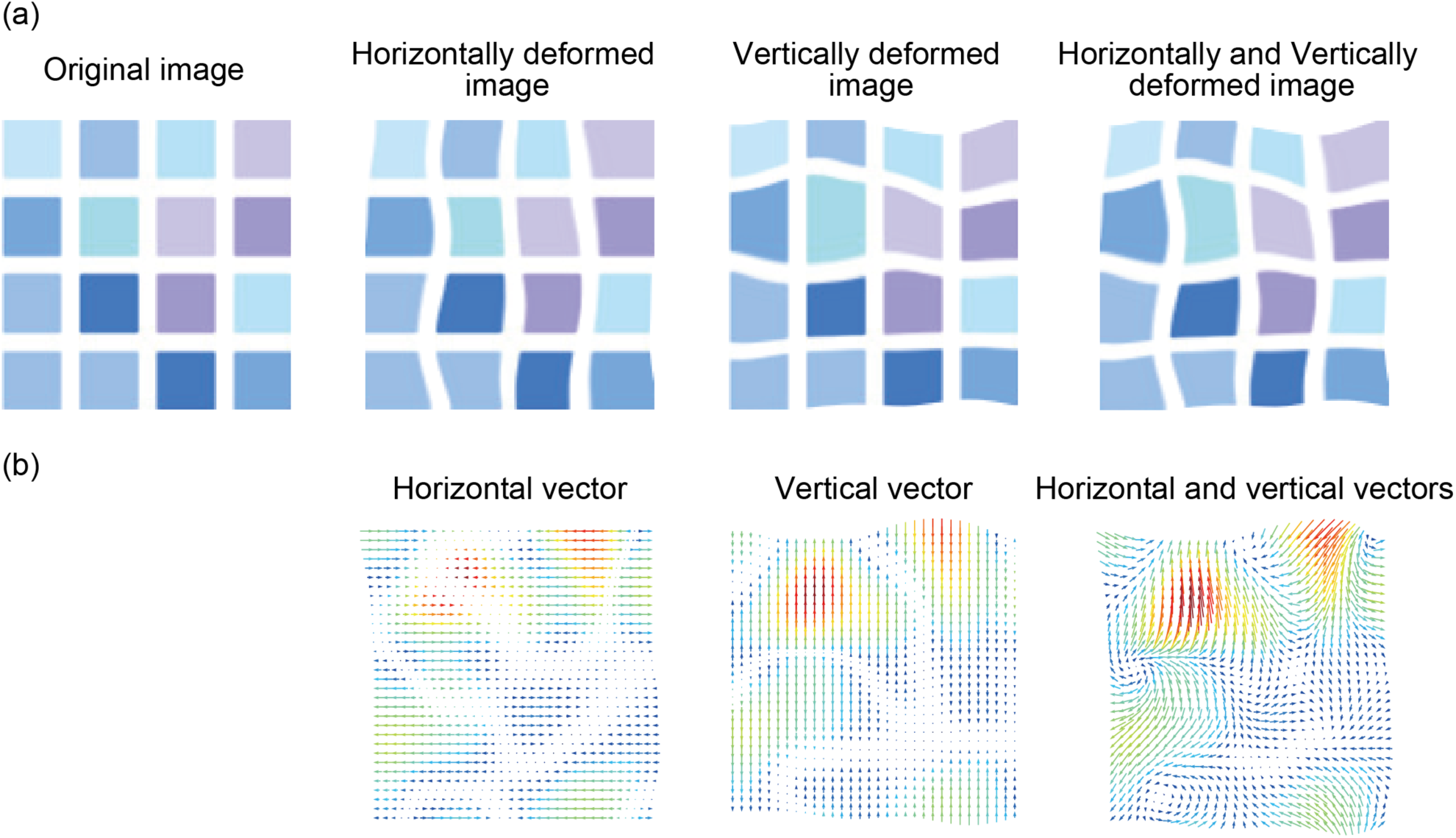
(a) Some examples of the appearance of an original image that is horizontally deformed, vertically deformed, and both horizontally and vertically deformed. (b) The deformation vector maps for horizontal deformation, vertical deformation, and both together.

There was also the possibility that the emergent global motion in a deformation pattern might be a cue to the direction discrimination of deformation flow. Studies [10] showed that shearing deformation spatially produced wavy patterns (Figure 2) and eventually generated unidirectional global motion on the basis of the intersection of constraint lines [11] when the amplitude of deformation was large. Thus, it was also predicted that the visual system determined the direction of deformation flow by using the global motion signals in image deformation.

**Figure 2.**
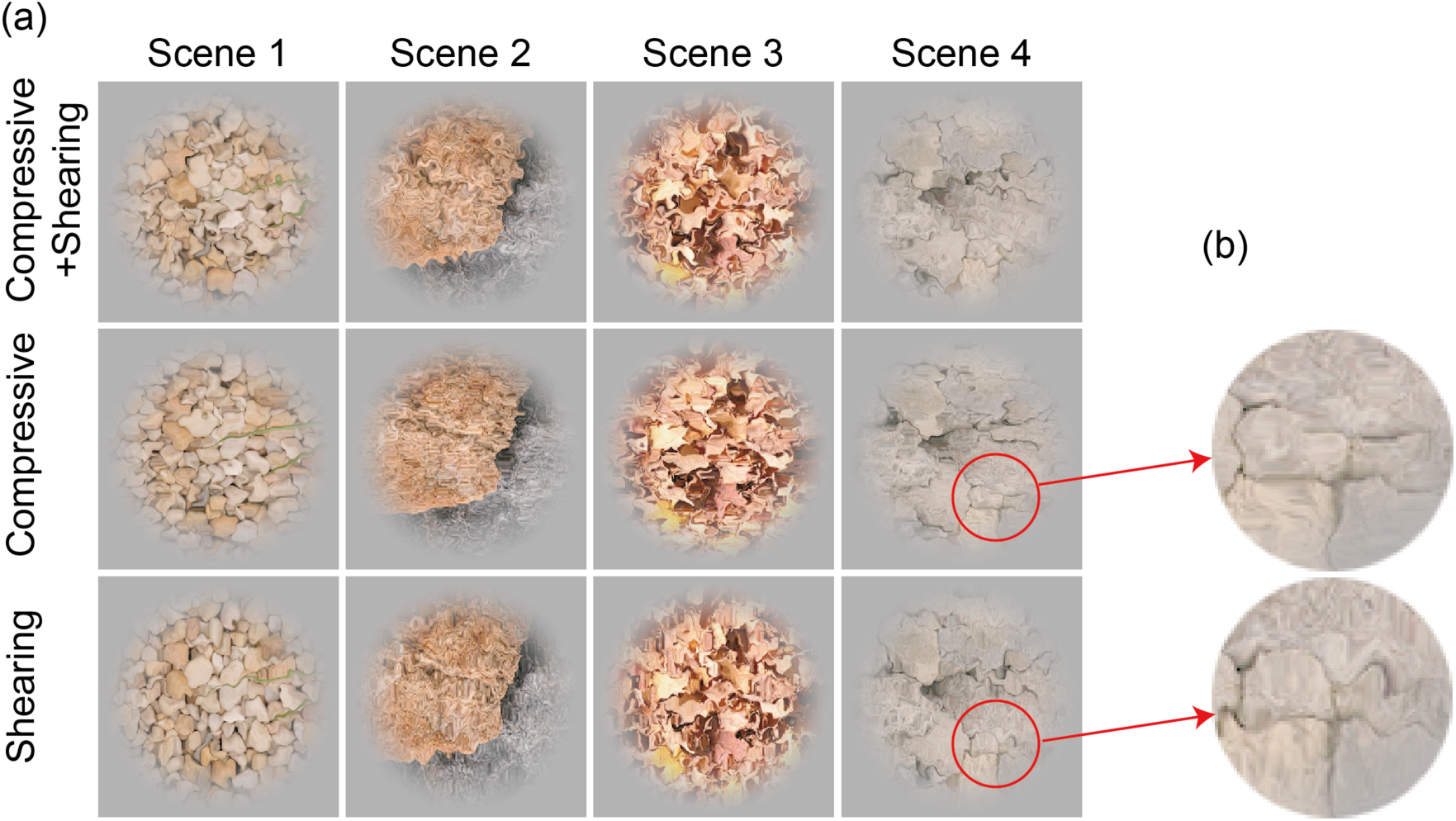
(a) Snapshots of stimulus clips as used in the experiments. We used four natural images as the background of stimuli. These snapshots were drawn from the stimulus clips in the 0.36 deg amplitude condition. (b) Enlarged part of snapshots of stimulus clips in the compressive and shearing conditions. The snapshot in the shearing condition contains the spatial wavy pattern.

The purpose of this study was two-fold. The first purpose was to psychophysically test how the human observers discriminated the direction of deformation flow. Manipulating the amplitude of image deformation, we checked how the amplitude influenced the discrimination performance. Also, we tested how the type of image deformation affected the performance. As described above, dynamic image deformation likely coming from transparent liquid flow consists of the compressive and shearing deformation, and the human visual system has different mechanisms for seeing compressive and shearing deformations [9]. It was thus worth testing whether the discrimination performance was affected differently by compressive and shearing deformations. The second purpose was to clarify what image motion cues were effective in the discrimination of deformation flow direction. By using a phase-based optical flow detection technique [12], which solves an aperture problem on the basis of the interaction of constraint lines, we investigated the relationship between the direction discrimination performance and the statistics of global motion. Based on the psychophysical and computational findings we discuss how human observers discriminate the direction of image deformation flow by using global motion cues in the deformation.

## Results

### Psychophysical experiment

In this experiment, we asked the observers to discriminate whether the transparent liquid in a stimulus clip (see Supplementary videos 1-3) translated leftward or rightward. In Figure 3 the proportion of trials in which the observer correctly reported the flow direction is plotted as a function of the amplitude of image deformation. Using the proportion, we conducted a two-way repeated measures ANOVA with deformation type and amplitude as within-subject factors. The main effect of deformation type was significant [*F*(2,12) = 115.08, p < .0001, *η*^*2*^_*p*_ = 0.95.] Multiple comparison tests showed that each deformation type was significantly different from the other (*p* < .0001). The main effect of amplitude was also significant [*F*(5,30) = 11.920, *p* < .0001, *η*^*2*^_*p*_ = 0.66]. Interaction between the two factors was also significant [F(10,60) = 8.211, *p* < .0001, *η*^*2*^_*p*_ = 0.58]. Further analysis [i.e., multiple comparison tests on the basis of the significant simple main effect (*p* < .0001)] showed that for each amplitude level, each deformation type was significantly different from the other (p < .0001) except for the 0.36 deg amplitude condition, which showed no significant difference between the compressive+shearing and shearing deformation conditions (*p* > .05). The analysis also showed that in the compressive+shearing deformation condition, the proportion in the 0.36 deg amplitude condition was significantly larger than the proportions in 0.06, 0.12, 0.18, and 0.24 deg amplitude conditions (*p* < .0001). Finally, for each deformation type, we also conducted a one-sample t-test with a Holm’s correction between the proportion of each amplitude condition and the chance level (0.5). For the compressive+shearing deformation, only the proportion of the 0.36 deg condition significantly deviated from 0.5 (*p* = .0022). For the compressive deformation, only the proportion of the 0.06 deg condition did not differ significantly from 0.5 (*p* = 0.55). For the shearing deformation, the proportions in all amplitude conditions differed significantly from 0.5 (*p* < .003).

**Figure 3.**
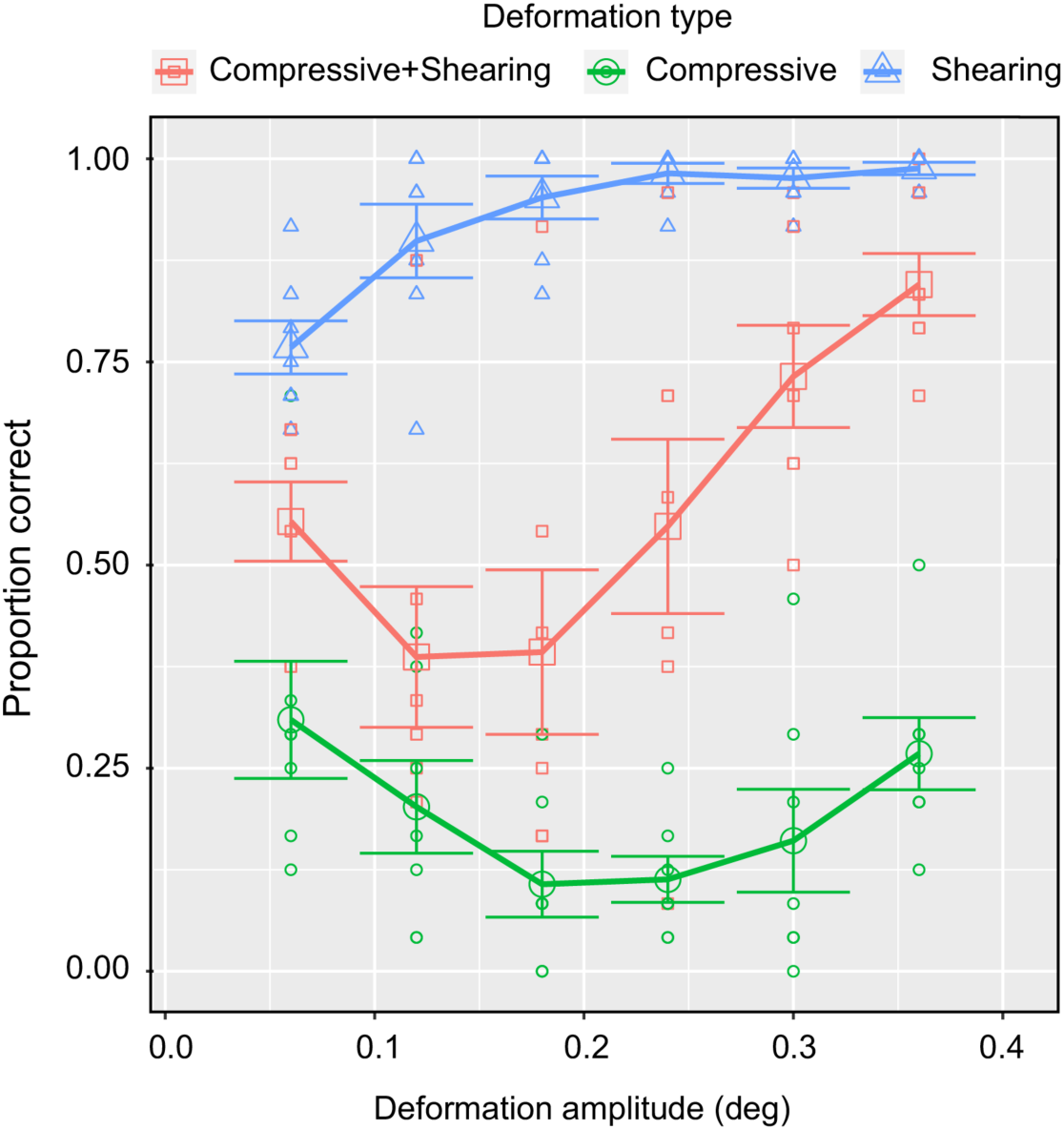
The result of the psychophysical experiment. Error bars denote ±1 standard errors of the mean (N = 7)

The results indicate that the observers could discriminate the direction of deformation flow when the image deformation amplitude was large enough, and also both compressive and shearing deformations were included in the stimulus clip. Moreover, we found an interesting effect of the deformation type on the perception of flow direction: the shearing deformation led to superior direction discrimination performance compared with the compressive and the compressive+shearing deformations. Importantly, the compressive deformation led to the perception of a flow direction opposite to the actual flow direction.

As a possible explanation of the shearing deformation result consistent with the previous study [10], emergent global motion signals on the basis of spatial wavy patterns served as a cue to the direction discrimination. In the compressive deformation case, however, the spatial wavy patterns were not more salient than the shearing deformation condition, possibly explaining why the discrimination performance was poor. Although it was not clear why the compressive deformation condition produced the perception of a flow direction opposite to the actual flow, we surmise that the compressive deformation also produced the peculiar global motion signals (or the first-order motion signals) that may have contributed to the perception of opposite flow direction. In the next section, we computationally analyze the image motion statistics of each deformation flow in stimulus clips and explore the relationship between discrimination performance and image motion statistics.

### Computational analysis

Image motion vectors of the stimulus clips were analyzed by using a phase-based optical flow detection algorithm [12]. The phase-based approach is akin to the spatiotemporal energy-based model for human motion vision [1]. Moreover, the algorithm solves an aperture problem [11,13] by using the interaction of constraint lines. Using the algorithm for each deformation type with each deformation amplitude, we calculated dense optical flow fields (128 × 128 cells) for all video frame pairs (29 pairs of 30 frames) in stimulus clips. Thus, we calculated 1,900,544 [128 (width) × 128 (height) × 29 (pairs of video frames) × 4 (types of background images)] motion vectors for each deformation type.

Based on the calculated motion vectors, we calculated several statistical indexes of horizontal motion vectors and plotted them in Figure 4 as a function of the amplitude of image deformation. We focused on horizontal motion vectors because the task of observers was to discriminate the horizontal direction (i.e., leftward or rightward) of deformation flow. Some important features can be identified. First, the mean values in the compressive+shearing condition were high when the amplitude was high. This pattern of change in the mean values is apparently consistent with the direction discrimination performance by the observers. In addition, the skewness in the compressive deformation condition was also apparently consistent with the direction discrimination performance.

**Figure 4.**
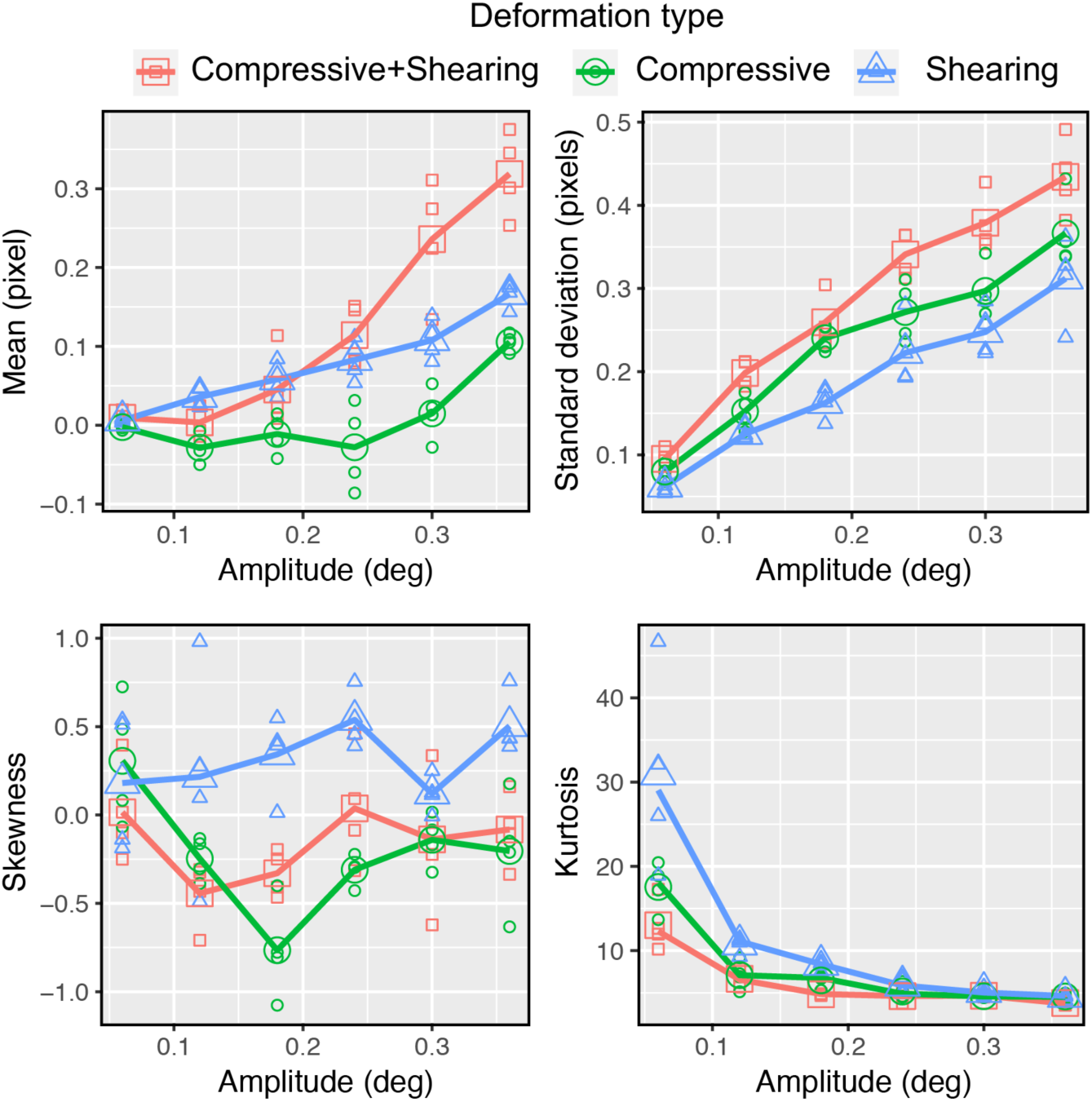
Statistical indices of horizontal vectors are plotted as a function of the amplitude of deformation. Top left: Mean values. Positive values denote motion direction in the actual flow direction. Top right: standard deviation, Bottom left: Skewness. Positive skewness indicates that the distribution has a longer tail in the direction of actual flow. Bottom right: Kurtosis.

To explore the contribution of the statistical indexes to the direction discrimination performance, by using the glmnet package in R [14], we conducted a Lasso regression [15] and found that mean and skewness contributed to the performance while standard deviation and kurtosis did not (Table 1). Figure 5 shows the outcome of the regression of a linear function to the discrimination performance as a function of the prediction using the coefficients in Lasso. The coefficient of determination (*r*^*2*^) was 0.69.

**Table 1.** The results of Lasso regression.

| **Statistics index** | **Coefficients** |
| --- | --- |
| (Intercept) | -1.142109e-16 |
| Mean | 2.971608e-01 |
| Standard deviation | - |
| Skewness | 5.537321e-01 |
| Kurtosis | - |

**Figure 5.**
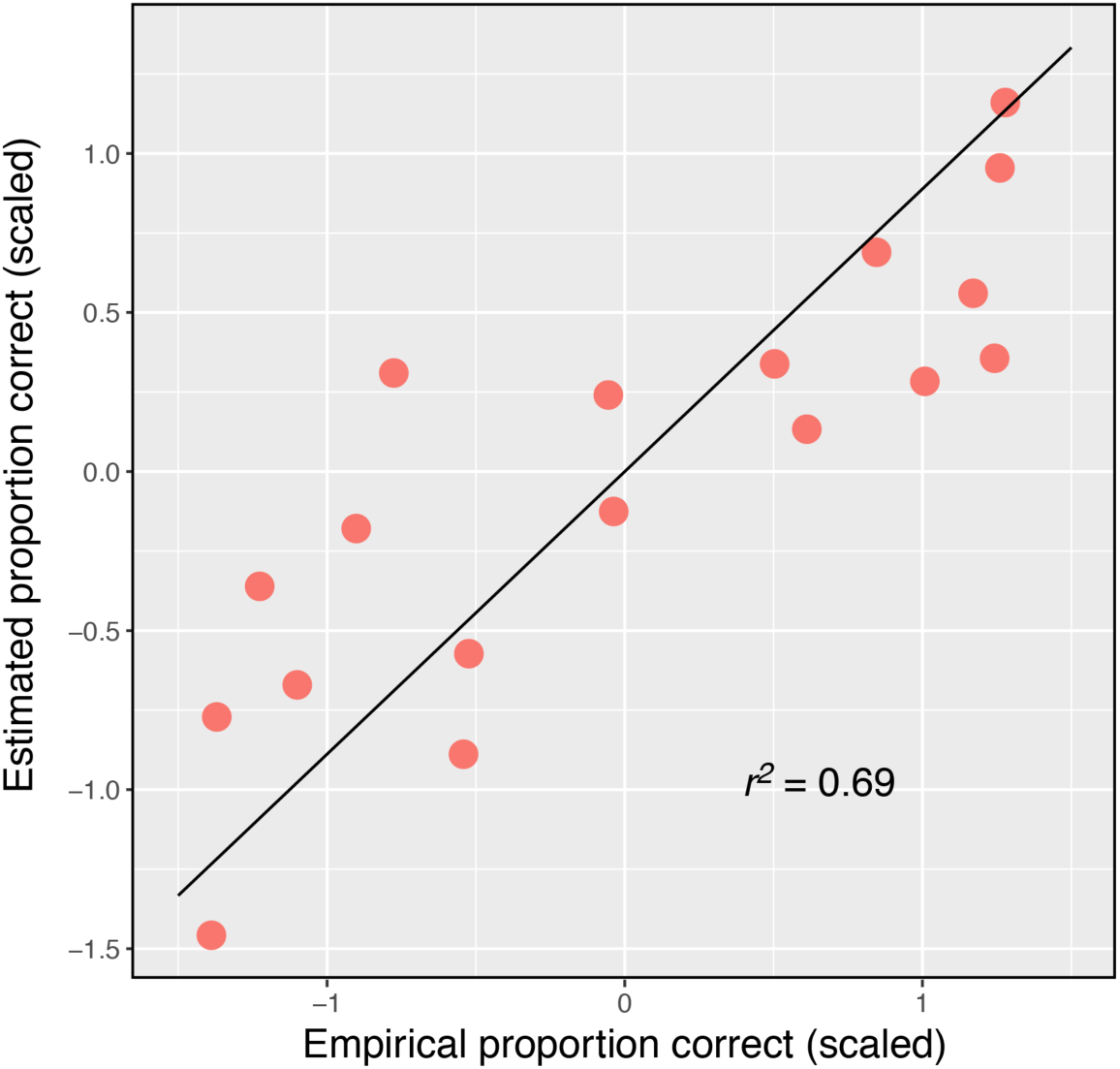
Estimated proportion correct that is based on Lasso regression is plotted as a function of empirical proportion correct (scaled).

Mean and skewness of the motion vectors were related to the bias of the direction of motion vectors, while standard deviation and kurtosis were not. This may be why only mean and skewness contributed to the prediction of the discrimination performance in Lasso regression. The results indicate that the biased global motion direction determines the perceived direction of deformation flow.

## General Discussion

We examined how human observers discriminated the direction of deformation flow. We found that when the deformation flow consisted of the compressive and shearing deformations, the observers could discriminate the direction of deformation flow only when the deformation amplitude was large. The discrimination performance was above the chance level irrespective of the deformation amplitude when the deformation flow consisted of the shearing deformation, while the observers consistently reported the flow direction opposite to the actual one when the deformation flow consisted of the compressive deformation. We tried to attribute the pattern of the discrimination performance to the statistical indexes of the horizontal motion vectors. Based on the Lasso regression, we found that the mean and skewness of the motion vectors significantly contributed to the prediction of the discrimination performance.

Because in our model the motion vectors were calculated on the basis of the intersection of constraint lines, the results indicate that the statistics of global motion are an effective descriptor of the performance. It is known that global motion can be integrated across space [16]. The integration mechanism may recruit the global motion having similar speeds across space and involves the determination of an overall flow direction.

Although both mean and skewness of the motion vectors were significant contributors to the prediction of the direction discrimination performance, the results do not always suggest that human observers can directly extract the statistical aspects of motion vectors. Specifically, it is unclear whether the visual system calculates the skewness of the motion vectors. With the stimuli we employed, when the skewness of the motion vectors was negative, the distribution had a longer tail in the motion direction opposite to an actual flow direction. That is, global motion vectors opposite to an actual flow direction tended to have a greater norm (i.e., speed) than global motion vectors consistent with the actual flow direction. The visual system may have employed the motion vectors with a greater speed as a cue to determine the flow direction. Although some previous studies have shown that image motion statistics could explain several aspects of dynamic material perception involving materials such as liquid [17] and fabric [18], the data do not always mean that the human visual system actually extracts the statistics. There is also the possibility that the brain extracts simpler motion information that is correlated with some sets of the statistical indexes. It will be necessary in future studies to address the question of whether the brain actually extracts the statistics of motion vectors using the more basic stimulus.

It is also possible that motion-defined motion contributed to the discrimination of flow direction. It is known that the visual system can extract the spatiotemporal flow of image signals that are defined by motion [19,20]. The motion-defined motion stimuli are not detected by conventional spatiotemporal energy detectors [1,20]. A sort of filter-rectify-filter scheme is often used to model the detection of higher-order motion such as motion-defined motion [21,22]. The algorithm we employed to calculate motion vectors is not sensitive to higher-order motion, while our stimuli likely contain motion-defined (or deformation-defined) flow. Because the Lasso regression on the basis of the statistical indexes of global motion explained 69% of the data, the remaining 31% might be partly explained by the contribution of the higher-order motion, which is left as an open issue for future studies.

We need to mention that transparent liquid flow in the natural world produces image flow other than the deformation-defined flow. For example, a specular flow is also caused by the liquid flow. It is known that the brain can use specular flow to identify surface material properties [23]. It is likely that in a more natural setting, the visual system uses specular flow as a cue to the discrimination of the liquid flow. However, given the motion vector pattern of the compressive deformation, it is likely that under some scenarios motion vectors of specular flow are not congruent with the motion vectors of global motion in the deformation. A previous study [24] has shown that the observer is not able to check the congruency between specular flow and deformation flow. Thus, given the incongruence of the motion vectors between specular flow and deformation flow, observers may use a specular flow as a cue to the discrimination of the liquid flow, while ignoring deformation-defined flow. On the other hand, specular flow is not always a reliable cue for direction discrimination because the appearance of specular flow is strongly affected by ambient illumination. In particular, on a cloudy day, specular reflection is not perceptually evident. Since the specular flow is not always employed as a cue in a fully reliable manner, the visual system may in parallel use both specular flow and deformation flow as cues to the discrimination of flow direction, following the principle of graceful degradation [25].

## Method

### Observers

Seven people (5 females and 2 males) participated in this experiment. Their mean age was 37.9 (SD: 7.1). All observers in this study reported having normal or corrected-to-normal visual acuity. They were recruited from outside the laboratory and received payment for their participation. Ethical approval for this study was obtained from the ethics committee at Nippon Telegraph and Telephone Corporation (Approval number: H28-008 by NTT Communication Science Laboratories Ethical Committee). The experiments were conducted according to principles that have their origin in the Helsinki Declaration. Written, informed consent was obtained from all observers in this study.

### Apparatus

Stimuli were presented on a 21-inch iMac (Apple Inc. USA) with a resolution of 1280 x 720 pixels and a refresh rate of 60 Hz. A colorimeter (Bm-5A, Topcon, Japan) was used to measure the luminance emitted from the display. A computer (iMac, Apple Inc., USA) controlled stimulus presentation, and data were collected with PsychoPy v1.83 [26-27].

### Stimuli

Stimuli were 30-frame video clips wherein a natural image sequentially deformed on the basis of translating deformation vector maps.

#### Generation of deformation vector map

The deformation vector map was a low-pass filtered version of the two-dimensional white noises (256 × 256 pixels, corresponding to 7.68 × deg). The cut-off spatial frequency of the low-pass filter was set to 2.08 cycles per degree (cpd) so that the spatial frequency of image deformation satisfied the stimulus condition to see transparent liquid [6]. Two deformation vector maps were used, one for horizontal deformation and the other for vertical deformation. In our stimuli, the deformation vector maps translated horizontally by 0.03 deg on each frame, so we called the horizontal deformation the compressive deformation and the vertical deformation the shearing deformation. The amplitude of deformation was one of the following six levels (0.06,0.12,0.18,0.24,0.30, and 0.36 deg). For example, in the case of the 0.36 deg deformation, the motion vector norm of the deformation ranged between −0.36 and 0.36 deg, uniformly distributed.

#### Image deformation

A natural image was sequentially deformed upon the generated deformation vector maps. As shown in Figure 2, four natural images were drawn from the McGill Calibrated Colour Image Database [28]. For image deformation, a conventional image warp method was used, as implemented in OpenCV (cv2.remap function). After the image deformation, each video frame was spatially windowed with a two-dimensional Tukey window so that the artifact of image deformation along the edges of the image was eliminated.

#### Making a stimulus clip

The deformed images were assembled so that their sequence comprised a 30-frame clip wherein the duration of each frame was 0.167 sec, so the duration of an entire clip was 0.5 sec.

### Procedure

Each observer was tested in a lit chamber. The observers sat 64 cm from the display. With each trial, a stimulus clip was presented for 0.5 seconds. The observer was asked to judge whether the static bar dynamically deformed or not after the disappearance of the clip. The judgment was registered by pressing one of the assigned keys. Each observer had four sessions, each consisting of 3 deformation types × 6 deformation amplitudes × 4 backgrounds × 2 repetitions. Within each session the order of trials was pseudo-randomized. Thus, each observer had 576 trials in total. It took 30–40 minutes for each observer to complete the entire set of four sessions.

## References

1. Adelson, E. H., & Bergen, J. R. (1985). Spatiotemporal energy models for the perception of motion. Journal of the Optical Society of America A, Optics and Image Science, 2(2), 284–299.

2. Nishida, S., Kawabe, T., Sawayama, M., & Fukiage, T. (2018). Motion Perception: From Detection to Interpretation. Annual Review of Vision Science, 4(1), 501–523. https://doi.org/10.1146/annurev-vision-091517-034328

3. Mather, G., & Cavanagh, P. (1989). Motion: The long and short of it. Spatial Vision, 4(2–3), 103–129. https://doi.org/10.1163/156856889X00077

4. Lu, Z.-L., Liu, C. Q., & Dosher, B. A. (2000). Attention mechanisms for multi-location first- and second-order motion perception. Vision Research, 40(2), 173–186. https://doi.org/10.1016/S0042-6989(99)00172-8

5. Ashida, H., Seiffert, A. E., & Osaka, N. (2001). Inefficient visual search for second-order motion. JOSA A, 18(9), 2255–2266. https://doi.org/10.1364/JOSAA.18.002255

6. Kawabe, T., Maruya, K., & Nishida, S. (2015). Perceptual transparency from image deformation. Proceedings of the National Academy of Sciences, 112(33), E4620–E4627. https://doi.org/10.1073/pnas.1500913112

7. Kawabe, T., & Kogovšek, R. (2017). Image deformation as a cue to material category judgment. Scientific Reports, 7, 44274. https://doi.org/10.1038/srep44274

8. Kawabe, T. (2018). Linear Motion Coverage as a Determinant of Transparent Liquid Perception. I-Perception, 9(6), 2041669518813375. https://doi.org/10.1177/2041669518813375

9. Nakayama, K., Silverman, G. H., MacLeod, D. I., & Mulligan, J. (1985). Sensitivity to shearing and compressive motion in random dots. Perception, 14(2), 225–238.

10. Nakayama, K. & Silverman, G. H. (1988). The aperture problem--I. Perception of nonrigidity and motion direction in translating sinusoidal lines. Vision Research, 28(6), 739–746. https://doi.org/10.1016/0042-6989(88)90052-1

11. Adelson, E. H., & Movshon, J. A. (1982). Phenomenal coherence of moving visual patterns. Nature, 300(5892), 523–525. https://doi.org/10.1038/300523a0

12. Gautama, T., & Van Hulle, M. M. (2002). A phase-based approach to the estimation of the optical flow field using spatial filtering. IEEE Transactions on Neural Networks, 13(5), 1127–1136. https://doi.org/10.1109/TNN.2002.1031944

13. Fennema, C. L., & Thompson, W. B. (1979). Velocity determination in scenes containing several moving objects. Computer Graphics and Image Processing, 9(4), 301–315. https://doi.org/10.1016/0146-664X(79)90097-2

14. Ihaka, R., and R. Gentleman. 1996. R: a language for data analysis and graphics. J. Comp. Graph. Stat. 5:299–314. Available via http://www.R-project.org.

15. Tibshirani, R. (1994). Regression Shrinkage and Selection Via the Lasso. Journal of the Royal Statistical Society, Series B, 58, 267–288.

16. Amano, K., Edwards, M., Badcock, D. R., & Nishida, S. (2009). Adaptive pooling of visual motion signals by the human visual system revealed with a novel multi-element stimulus. Journal of Vision, 9(3), 4. https://doi.org/10.1167/9.3.4

17. Kawabe, T., Maruya, K., Fleming, R. W., & Nishida, S. (2015). Seeing liquids from visual motion. Vision Research, 109, Part B, 125–138. https://doi.org/10.1016/j.visres.2014.07.003

18. Bi, W., Jin, P., Nienborg, H., & Xiao, B. (2019). Manipulating patterns of dynamic deformation elicits the impression of cloth with varying stiffness. Journal of Vision, 19(5), 18–18. https://doi.org/10.1167/19.5.18

19. Zanker, J. M. (1993). Theta motion: a paradoxical stimulus to explore higher order motion extraction. Vision Research, 33(4), 553–569. https://doi.org/10.1016/0042-6989(93)90258-X

20. Maruya, K., Mugishima, Y., & Sato, T. (2003). Reversed-phi perception with motion-defined motion stimuli. Vision Research, 43(24), 2517–2526. https://doi.org/10.1016/S0042-6989(03)00438-3

21. Chubb, C., & Sperling, G. (1989). Two motion perception mechanisms revealed through distance-driven reversal of apparent motion. Proceedings of the National Academy of Sciences, 86(8), 2985–2989. https://doi.org/10.1073/pnas.86.8.2985

22. Lu, Z.-L., & Sperling, G. (1995). The functional architecture of human visual motion perception. Vision Research, 35(19), 2697–2722. https://doi.org/10.1016/0042-6989(95)00025-U

23. Doerschner, K., Fleming, R. W., Yilmaz, O., Schrater, P. R., Hartung, B., & Kersten, D. (2011). Visual Motion and the Perception of Surface Material. Current Biology, 21(23), 2010–2016. https://doi.org/10.1016/j.cub.2011.10.036

24. Kawabe, T., & Nishida, S. (2015). Seeing transparent liquids from refraction-based image deformation and specular reflection. Journal of Vision, 15(12), 935–935. https://doi.org/10.1167/15.12.935

25. Marr, D. (1975). Early processing of visual information. AI Memos, No. 340.

26. Peirce, J. W. (2007). PsychoPy—Psychophysics software in Python. Journal of Neuroscience Methods, 162(1–2), 8–13. https://doi.org/10.1016/j.jneumeth.2006.11.017

27. Peirce, J. W. (2009). Generating stimuli for neuroscience using PsychoPy. Frontiers in Neuroinformatics, 2, 10. https://doi.org/10.3389/neuro.11.010.2008

28. Olmos, A., & Kingdom, F. A. A. (2004). A biologically inspired algorithm for the recovery of shading and reflectance images. Perception, 33(12), 1463–1473.

